# Acquisition of drug resistance in basal cell nevus syndrome tumors through basal to squamous cell carcinoma transition

**DOI:** 10.1101/2023.07.26.550719

**Authors:** Anna R. Jussila, Daniel Haensel, Sadhana Gaddam, Anthony E. Oro

## Abstract

While basal cell carcinomas (BCCs) arise from ectopic hedgehog pathway activation and can be treated with pathway inhibitors, sporadic BCCs display high resistance rates while tumors arising in Gorlin syndrome patients with germline Patched (*PTCH1*) mutations are uniformly suppressed by inhibitor therapy. In rare cases, Gorlin syndrome patients on long-term inhibitor therapy will develop individual resistant tumor clones that rapidly progress, but the basis of this resistance remains unstudied. Here we report a case of an SMO^i^-resistant tumor arising in a Gorlin patient on suppressive SMO^i^ for nearly a decade. Using a combination of multi-omics and spatial transcriptomics, we define the tumor populations at the cellular and tissue level to conclude that Gorlin tumors can develop resistance to SMO^i^ through the previously described basal to squamous cell carcinoma transition (BST). Intriguingly, through spatial whole exome genomic analysis, we nominate PCYT2, ETNK1, and the phosphatidylethanolamine biosynthetic pathway as novel genetic suppressors of BST resistance. These observations provide a general framework for studying tumor evolution and provide important clinical insight into mechanisms of resistance to SMO^i^ for not only Gorlin syndrome but sporadic BCCs as well.

## INTRODUCTION

Targeted therapies show efficacy in treating many different cancers, but it has become clear that factors such as tumor heterogeneity reduce their overall effectiveness. Heterogeneity comes from intrinsic resistance from pre-existing tumor populations as well as a dynamic intrinsic cellular plasticity, allowing tumor cells to toggle to differential phenotypic states when challenged with therapy or the accumulation of driver mutations. A clearer understanding of the discrete cellular populations capable of phenotype switching, the barriers to such switching, and the overall molecular genetic and epigenetic drivers that facilitate these transitions will aid in constructing precision cancer treatment strategies (Boumahdi and de Sauvage, 2020).

With well-defined tumor lineage relationships, a general understanding of many of their key molecular drivers, and the ability to perform serial biopsies, basal cell carcinomas (BCCs) and squamous cell carcinomas (SCCs) serve as ideal models to deconvolute the dynamic tumor heterogeneity and its impact on tumor growth and drug resistance (Atwood et al., 2015; Sanchez-Danes and Blanpain, 2018). The Hedgehog (HH) signaling pathway is the key molecular driver of sporadic BCCs, with known driver mutations occurring in the key signaling components such as Patched (*PTCH1*) or the G-protein coupled receptor Smoothened (*SMO*) (Oro et al., 1997). SMO inhibitors (SMO^i^) have proven extremely useful as a primary therapy for those patients with advanced tumors that are not surgically resectable (Sekulic et al., 2012). Unfortunately, approximately 60% of patients are initially resistant, and a further 20% of patients subsequently acquire resistance to SMO^i^, indicating that deconvolution of the various resistance mechanisms is still needed (Atwood *et al.*, 2015).

In contrast to patients with sporadic BCCs, patients with Gorlin syndrome that possess a germline *PTCH1* mutation develop BCCs all over their body from an early age and throughout their lifetime (Tang et al., 2012). Like sporadic BCCs, Gorlin BCC tumors are driven by the upregulation of HH signaling and SMO^i^ therapy effectively suppresses Gorlin patient BCCs with an extremely low resistance rate (Tang *et al.*, 2012). While SMO^i^ therapy reduces Gorlin patients’ BCC tumor burden, small tumor populations persist at the site of the original tumor, and if patients are taken off therapy, these persister populations reemerge as tumors in the same location (Tang *et al.*, 2012). Due to low mutational burden and intact DNA repair systems, tumor evolution remains low and can be continually suppressed with re-initiation of the SMO^i^ therapy, facilitating drug holidays for users (Chiang et al., 2018). The basis of this surprising lack of tumor progression and the rarity of drug-resistant BCCs in Gorlin patients remains puzzling but provides a clinically important question to address.

Recent work from our lab aimed at elucidating resistance mechanisms associated with locally advanced or metastatic resistant BCCs (rBCCs) identified a novel resistance mechanism called the basal to squamous cell carcinoma transition (BST), which is characterized by the development of well-differentiated SCC-like histological features within sporadic BCCs in response to SMO^i^ treatment (Haensel et al., 2022; Kuonen et al., 2021). We have previously shown that these SCC-like populations exist in sporadic naive BCCs (nBCCs), are marked by the surface marker Lymphocyte Antigen 6 Family Member D (LY6D), and expand upon SMO^i^ treatment (Haensel *et al.*, 2022). *LY6D*^+^ cells do not express HH-related genes like *GLI1*, indicative of HH-independent growth and consequently SMO^i^ resistance (Haensel *et al.*, 2022). Interestingly, while sporadic BCCs frequently convert to well-differentiated SCCs with high *LY6D* expression, Gorlin patient persister BCCs, although they express high levels of *LY6D* during SMO^i^ treatment, generally remain small and do not grow out as converted well-differentiated SCCs. Furthermore, it appears that after SMO^i^ removal, these persister tumors can revert to an *LY6D*^-^ and SMO^i^-sensitive state (Haensel *et al.*, 2022).

In this work, we interrogate a distinct suppression-resistant Gorlin BCC that arose within a previously drug-suppressed BCC. In contrast to typical Gorlin BCCs, we find that this tumor has grown out as an SCC-like tumor, with characteristics suggestive of a BST-associated resistance mechanism. We take advantage of new spatial transcriptomic technologies, integrating with single-cell RNA sequencing (scRNA-Seq) to provide a detailed spatial analysis of tumor epithelial dynamics associated with BST. We find that at the gene expression level, the tumor populations from the SMO^i^-treated Gorlin tumor transition towards an HH-independent state, resembling BST-like populations seen in sporadic and resistant BCCs. Through spatial whole exome sequencing (WES) genomics, we show that the BCC- and SCC-like portions of the tumor are indeed related by lineage and that the SCC-like portion has two mutations within the phosphatidylethanolamine (PE) synthesis pathway, *PCYT2, and ETNK1.* Inhibition of the PE synthesis pathway causes gene expression changes consistent with BST in BCC cells, nominating the PE synthesis pathway as a novel driver of BST.

## RESULTS

### Identification of BST in resistant basal cell nevus syndrome (Gorlin Syndrome) tumor

The 56-year-old Gorlin patient, diagnosed at age 7 due to odontogenic jaw cysts but without medulloblastomas, had been on tumor-suppressive doses of SMO^i^ for over 10 years and managed the tumor burden with preventative agents such as photodynamic therapy and topical imiquimod without surgical intervention. Despite this history, a focal rapidly growing 2 cm tumor suddenly appeared while on therapy and was removed from the left neck (Figure 1a). Histological examination found that the tumor displayed a spectrum of BCC- and SCC-like phenotypes (Figure 1b). Intrigued by the apparent squamous populations within the tumor, we checked for markers of BST and found that the SCC-like portion expressed high levels of *LY6D* (Figure 1c). In contrast, the BCC-like portion of the tumor expressed high levels of *GLI1* (Figure 1c). There were a few *GLI1*^+^ cells within the SCC-like tumor epithelium, typically at the stromal interface but none were detected in more centralized portions of the tumor nodules (Figure 1c). Overall, these findings suggest that suppression-resistant Gorlin tumors escape through the *LY6D* BST plasticity pathway. Although we have previously shown that Gorlin persister populations express low levels of *LY6D* when treated with SMO^i^, extensive growth of SCC-like *LY6D*-expressing tumor populations during SMO^i^ treatment has not previously been observed and has critical implications for the treatment of Gorlin tumors as well as sporadic BCC (Haensel *et al.*, 2022).

**Figure 1:**
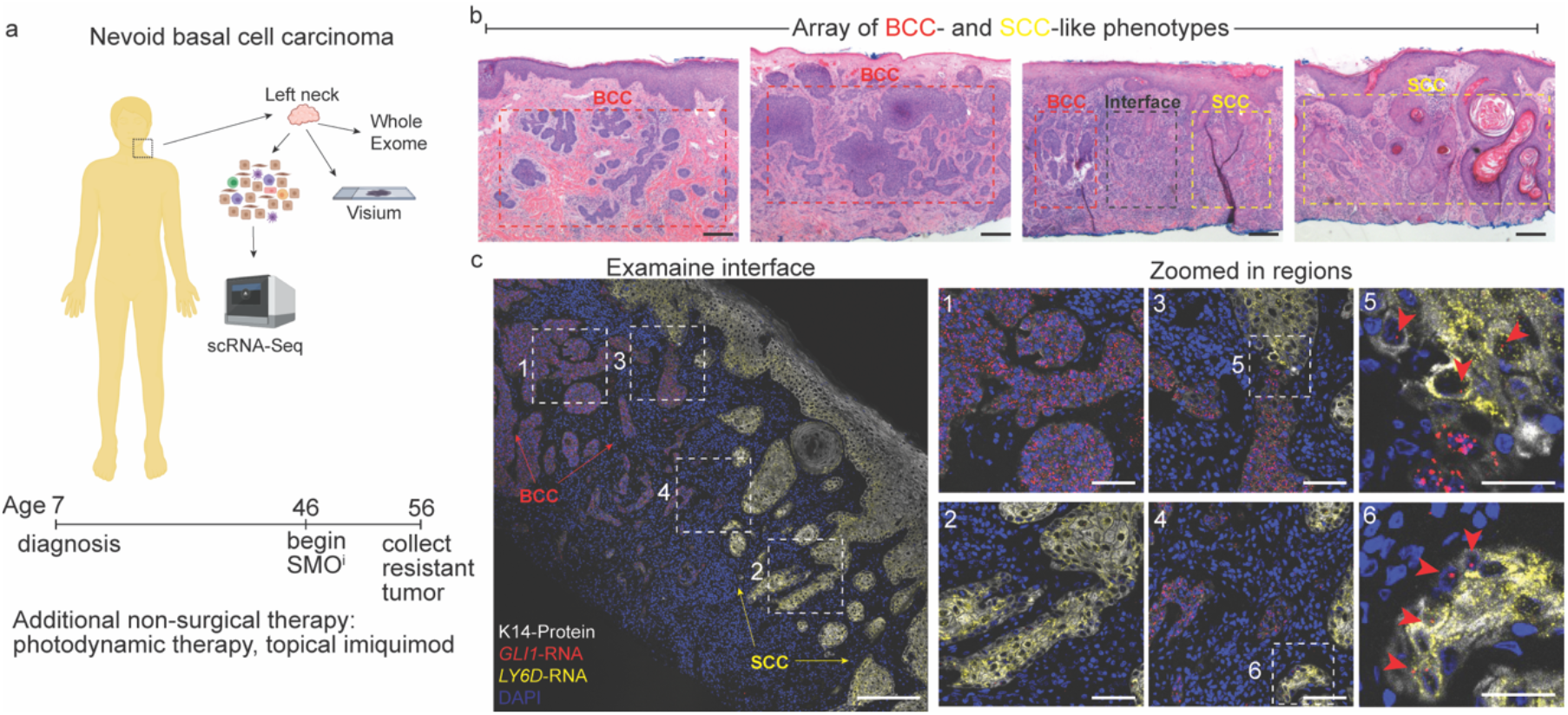
Identification of BST in resistant basal cell nevus syndrome (Gorlin Syndrome) a. Schematic diagram of extracted tumor locations and subsequent downstream processing and analysis with timeline describing treatment overview. b. H/E of the tumor from the left neck with distinct BCC- and SCC-like features annotated. Scale bars = 50μm. c. RNAScope analysis at the interface between the BCC- and SCC-like regions with *GLI1* (red) and *LY6D* (yellow). Co-staining was done with K14 protein (white) and Hoechst (blue). Scale bars = 200μm for large image, 50μm for regions 1-4, and 25μm for regions 5 and 6.

### Coupled scRNA-Seq and spatial transcriptomics to profile resistant Gorlin tumor reveals varied tumor organization

To comprehensively profile this resistant Gorlin tumor, we conducted a combination of scRNA-Seq and spatial transcriptomics on the same sample (Figure 1a). Using scRNA-Seq, we were able to identify 17 different clusters of cells comprising a combination of different cell types and cell states (Figure 2a, Supplementary Figure S1a, and Supplementary Table S1). Using various differential and known marker genes associated with major cell types, we were able to identify tumor (*KRT14^+^GLI1^+^*) and normal epithelial populations (*KRT14^+^GLI1^-^*) as well as stromal components including fibroblasts (*COL1A2^+^*), endothelial cells (*PECAM1^+^*), melanocytes (*PMEL^+^*), and different immune cell populations such as myeloid cells (*PTPRC^+^CD68^+^*) and T cells (*PTPRC^+^CD3E^+^*) (Figure 2a and b) (Haensel *et al.*, 2022; Haensel et al., 2020).

**Figure 2:**
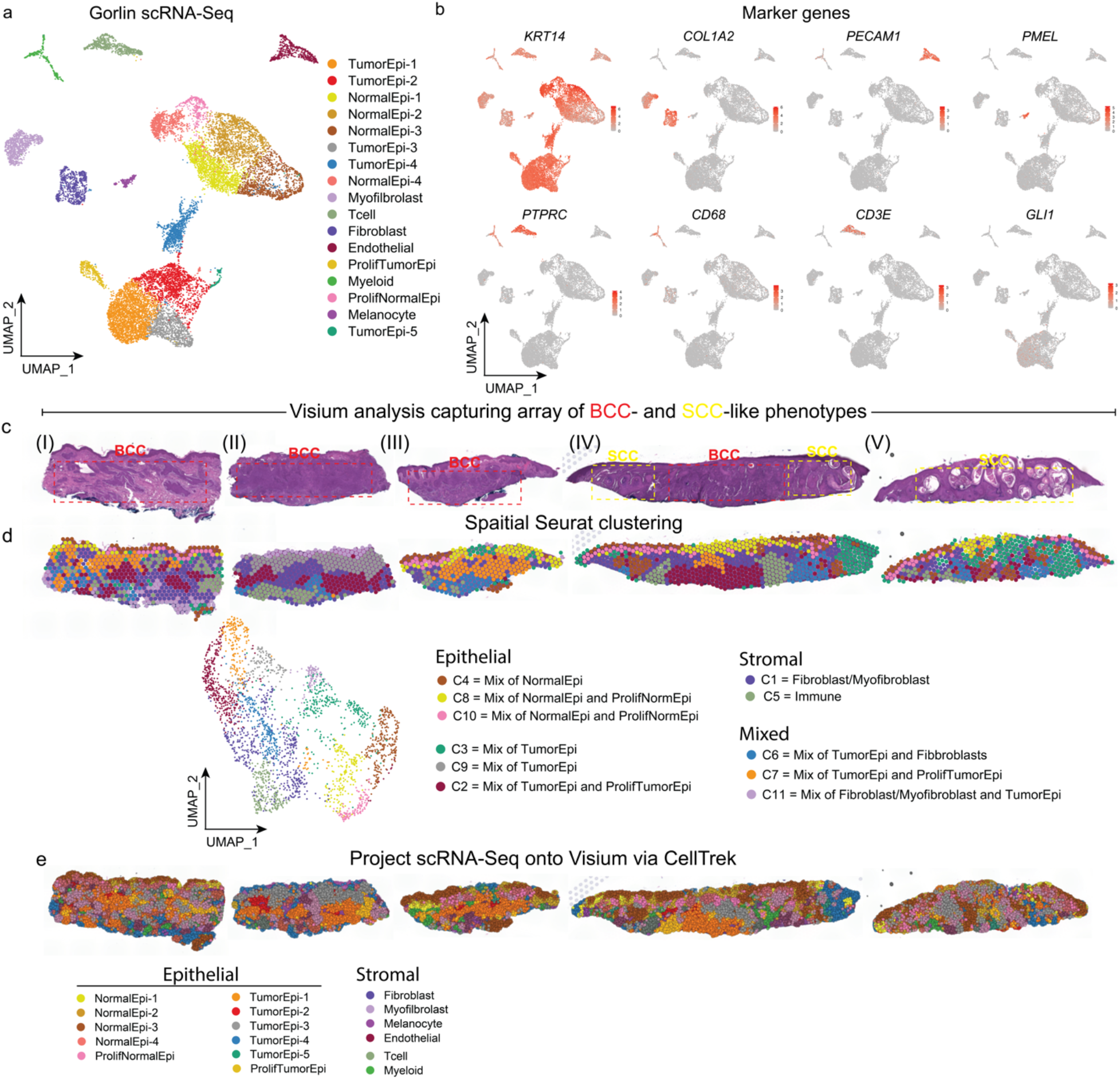
Coupled scRNA-Seq and spatial transcriptomics to profile resistant Gorlin tumor reveals varied tumor organization. a. UMAP plot of scRNA-Seq data of all cells (both tumor and adjacent normal tissue) from the Gorlin sample containing 17 clusters. b. Feature plots highlighting key marker genes used to identify the different clusters in (a). c. Overview of the Visium analysis containing the five different captured tissue sections (I- V) including both the H&E, which highlights the histologically distinct BCC- and SCC- like regions captured. d. UMAP plot of the merged spatial Seurat analysis from the five different sections. e. Projection of scRNA-Seq data onto Visium analysis of tissue sections (I)-(V) using CellTrek.

We further used the 10x Visum platform for spatial transcriptomics, capturing regions (I) through (V) that span the histologically distinct tumor regions defined by BST (Figure 2c and Supplementary Figure S1b). Region (I) has a nodular BCC-like appearance characterized by epithelial cells with little cytoplasm arranged in well-defined palisades (Figure 2c). Regions (II) and (III) have less well-defined tumor nodules and a more infiltrative BCC-like appearance (Figure 2c). Region (IV) has a combination of infiltrative BCC-like regions and well-differentiated SCC-like regions, characterized by eosinophilic cytoplasm among tumor cells and increased keratinization (Figure 2c). Region (V) is entirely SCC-like in appearance (Figure 2c). By performing spatial dimensional reduction, we identified 11 clusters of spatial transcriptomic spots and determined their composition manually by scoring for the marker genes from our previously labeled scRNA-Seq clusters (Figure 2a, d, Supplementary Figure S1c, d, and Supplementary Table S2).

As expected, given the capture area size of the Visium platform, some of the clusters had hybrid gene expression patterns, sharing both epithelial and stromal gene expression signatures, likely reflecting the simultaneous capture of different cell types (Figure 2d and Supplementary Figure S1d). To further deconvolute the composition of multi-cell capture spots, we used the program CellTrek, which utilizes our scRNA-Seq data in combination with a random forest classifier to superimpose scRNA-Seq data onto the spatial Visum analysis (Figure 2e) (Wei et al., 2022). We first confirmed CellTrek’s ability to accurately map the normal epidermal cells and found that the normal epithelial cells were for the most part placed along the epidermis (Supplementary Figure S2a). Continuing to use the histology as a reference, CellTrek largely accurately superimposes tumor epithelial cells on regions that are recognizably tumor by histological examination and stromal cells in non-tumor regions (Figure 2e and Supplemental Figure S2b). In addition, using the histology to validate CellTrek predictions, we also utilized the marker gene signatures from the different epithelial clusters of our scRNA-Seq (Figure 2e, Supplementary Figure 2b, Supplementary Figure 3b and c). Overall, Seurat feature scoring largely agreed with the CellTrek predictions and together, they can be used to overcome the multiple cell capture variable associated with Visium.

### Tumor epithelial populations occupy distinct regions within the Gorlin tumor

With the spatial locations established, we next investigated the spatial distribution of tumor epithelial cells in relation to the BCC- and SCC-like histological features of the tumor. Using our previously identified BST markers, we confirmed the spatial distribution of *LY6D*, *LYPD3*, and *TACSTD2* along with the BCC-associated marker, *GLI1* within the different tissue sections (Figure 3a and Supplementary Figure S1c) (Haensel *et al.*, 2022). In section (I), the *GLI1* was restricted to the BCC-like tumor, and the *LY6D* was restricted to the normal epidermis (Figure 3a). In section (V), which has exclusively SCC-like features, there was extensive *LY6D* and low *GLI1* expression (Figure 3a). The most striking observations are in section (IV) which has distinct BCC- and SCC-like regions (Figure 3a). In this section, there is little overlap between regions of high *GLI1* and *LY6D* expression (Figure 3a). Overall, this data suggests that Gorlin patients can develop resistance to SMO^i^ by mechanisms like BST based on the expression of associated marker genes.

**Figure 3:**
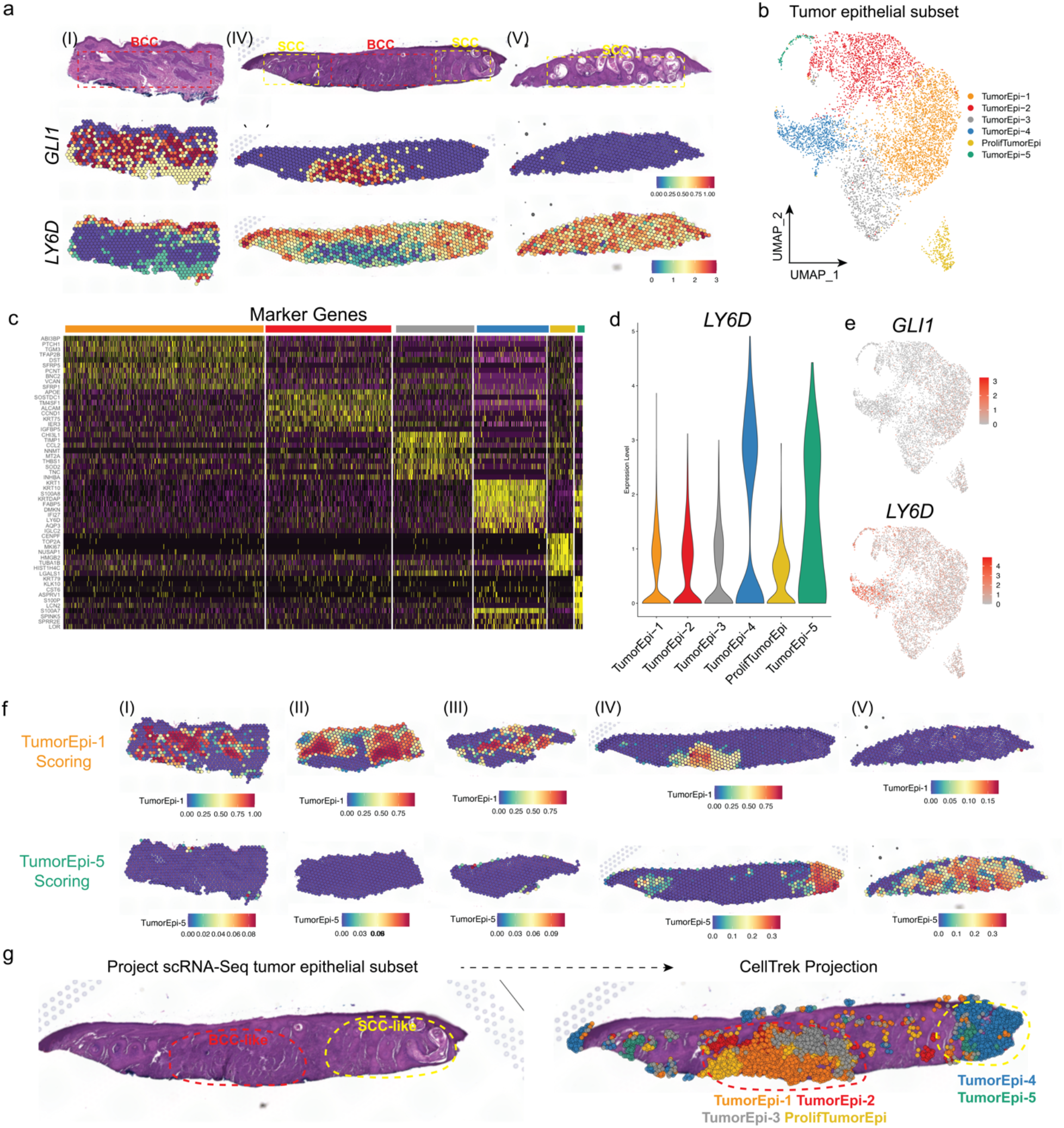
Spatial analysis and characterization of Gorlin tumor epithelial populations. a. Spatial feature plots of *GLI1* and *LY6D* for tissue sections (I), (IV), and (V). b. UMAP plot of re-clustered scRNA-Seq data of the six tumor epithelial clusters. c. Heatmap of marker genes associated with each of the six tumor epithelial clusters from (a). d. Violin plot of *LY6D* in each of the six tumor epithelial clusters from (a). e. Spatial feature plots of the tumor epithelial clusters for *GLI1* and *LY6D*. f. SpatialSeurat gene scoring for the TumorEpi-1 and TumorEpi-5 populations on all of the various tissue sections. g. CellTrek projection of the different scRNA-Seq tumor epithelial populations onto Visium analysis of tissue section (IV).

To further profile the tumor epithelium and to investigate BST in this Gorlin tumor more comprehensively, we subsetted and re-clustered the tumor epithelium from the scRNA-Seq, identifying 6 main clusters (Figure 3b and Supplementary Table S3). We identified the various marker genes and associated Gene Ontology (GO) terms associated with the various clusters (Figure 3c and Supplementary Figure S3a). Like our analysis in sporadic BCCs, we found that these differential tumor epithelial states were driven by GO terms associated with various differentiation and proliferation states (Supplementary Figure S3a). At the gene expression level, it appeared that TumorEpi-4 and TumorEpi-5 had the highest levels of *LY6D* and corresponding low levels of *GLI1* (Figure 3d, e).

To next identify where these different populations spatially reside within the tissue, we used a combination of spatial scoring and CellTrek predictions. Spatial scoring found that TumorEpi-1 was spatially localized to the BCC regions of sections (I)-(IV) (Figure 3f and Supplementary S3b).

Subsequent spatial feature scoring for TumorEpi-4 and TumorEpi-5 predicted that these cells were spatially localized to the SCC regions of sections (IV) and (V) (Figure 3f and Supplementary Figure S3b). For our CellTrek analysis, we focused on section (IV), which displayed both BCC- and SCC-like features. Like the spatial scoring, CellTrek predicted that TumorEpi-1, TumorEpi- 2, TumorEpi-3, and ProlifTumorEpi were localized to the BCC region while TumorEpi-4 and TumorEpi-5 were spatially localized to the SCC regions (Figure 3g and Supplemental Figure S3c). These data confirm that tumor epithelial populations differ between the regions of the tumor that display BCC-like features from those that display well-differentiated SCC-like features.

### Contextualization of BST across Gorlin, nBCCs, rBCCs, and SCCs

To better categorize these different tumor epithelial clusters and contextualize their placement along the BST spectrum, we used a combination of our previous analysis from naive non-drug treated sporadic BCCs, resistant BCCs, and existing data from SCCs (Haensel *et al.*, 2022; Ji et al., 2020; Yao et al., 2020; Yost et al., 2019). In our previous analysis of sporadic BCCs, we identified a particular cluster of cells from our scRNA-Seq analysis, which was enriched for BST- associated markers such as *LY6D* (which we referred to as Cluster 6/C6 in the original publication) (Haensel *et al.*, 2022). Taking the marker genes associated with this C6, we found that TumorEpi- 5 from our Gorlin analysis scored the highest for this gene signature (Figure 4a). In line with this enhanced scoring for this cell state, TumorEpi-5 also scored high for the ‘Persister’ population of cells, previously shown to not be responsive to SMO^i^ (Figure 4b) (Biehs et al., 2018). Our analysis would suggest that TumorEpi-5 shares similarities with the BST-associated populations we had identified in sporadic BCCs, but what is not clear is where exactly this population falls along the BST spectrum.

**Figure 4:**
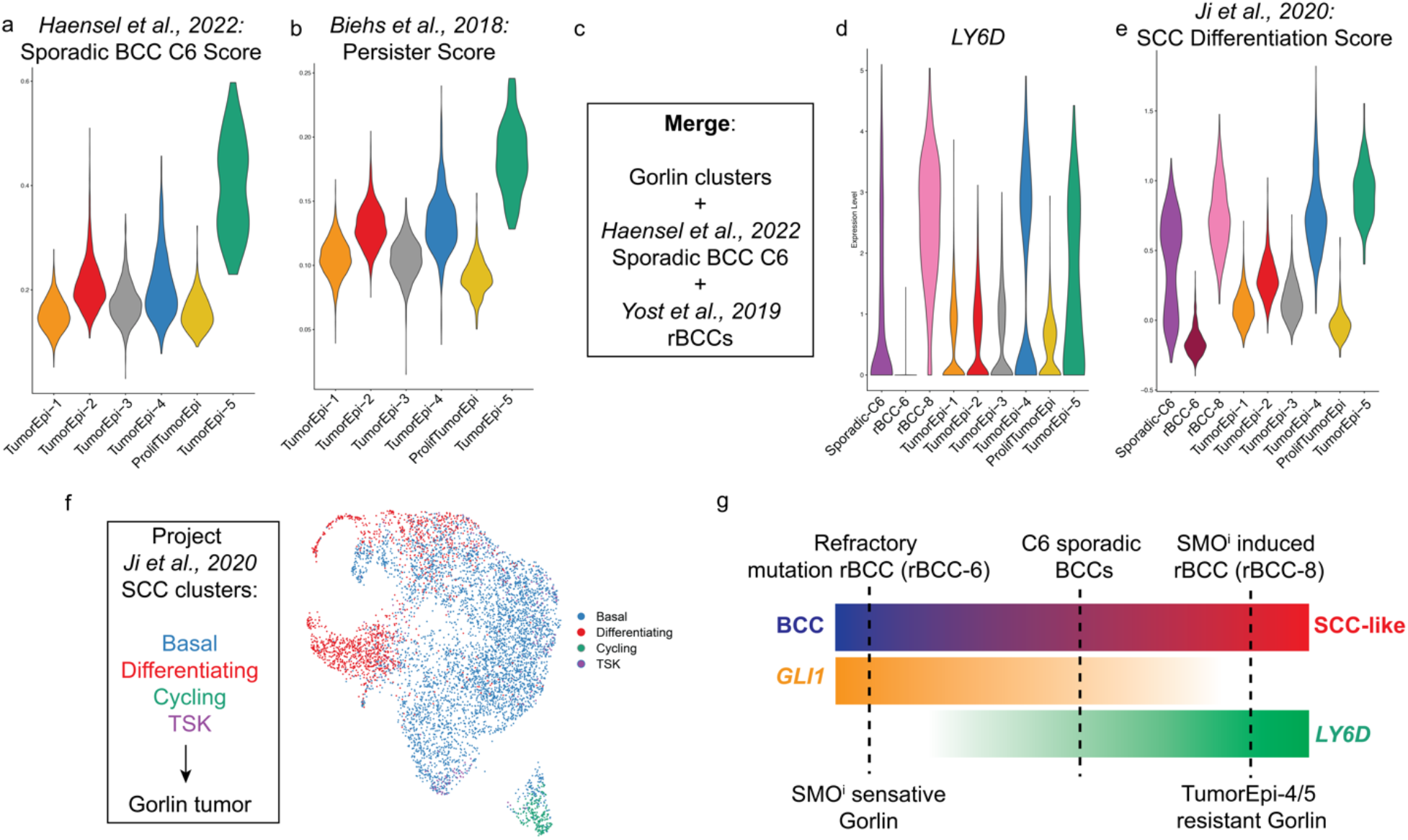
Contextualization of BST across Gorlin, nBCCs, rBCCs, and SCCs. a. Violin plot of Sporadic BCC C6 gene scoring for the six tumor epithelial clusters from b. Violin plot of Persister scoring for the six tumor epithelial clusters from (a). c. Merging strategy for different tumor epithelial subsets. d. Violin plot of *LY6D* in the merged tumor epithelial subset from (i). e. Violin plot of the SCC Differentiation score for the merged tumor epithelial subset from (i). f. Projection of SCC-associated cell states onto the Gorlin tumor epithelial subset from (a). g. Diagram of the BST spectrum and predicted placement of the various tumor types.

To address this, we directly compared the clusters from this Gorlin tumor to our previously discussed Cluster 6 from sporadic BCCs (Sporadic-C6) along with two different resistant BCC samples that we had previously analyzed; one that has BST-phenotypes and expresses *LY6D* (rBCC-8) and one that has no *LY6D* expression and a known gene mutation that allows it to evade SMO^i^ (rBCC-6) (Figure 4d) (Haensel *et al.*, 2022; Yost *et al.*, 2019). We had previously shown that tumors that have SMO-activating mutations, such as rBCC-6, do not have any *LY6D* expression and do not appear to utilize BST as a resistance mechanism (Haensel *et al.*, 2022). Comparing all these groups simultaneously, it appears that TumorEpi-4 and TumorEpi-5 have more substantial levels of *LY6D* than rBCC-6, such that their expression level is comparable to that of rBCC-8 and Sporadic-C6 (Figure 4d). To further contextualize these populations, we next used well-defined marker gene lists to define populations of cells from well-differentiated SCCs (Ji *et al.*, 2020). We found that TumorEpi-4 and TumorEpi-5 had very similar scoring to rBCC-8 (Figure 4e). This was further validated by projecting gene signatures for the different previously identified SCC-associated populations onto our Gorlin dataset (Figure 4f) (Ji *et al.*, 2020). Overall, this analysis suggests that both TumorEpi-4 and TumorEpi-5 appear to have progressed more along the BST spectrum compared to our Sporadic-C6 cluster from naïve BCCs, more closely resembling rBCC-8 (Figure 4g).

### Squamous regions within the Gorlin tumor are of BCC origin

With the resistant Gorlin tumor placed along the BST spectrum through Monocle analysis (Supplemental Figure S4a), we endeavored to demonstrate that the BCC- and SCC-like portions are related by lineage, confirming BST (in contrast to distinct coincidental BCC and SCC). In BSCs, whole exome sequencing (WES) has shown that the SCC-like portions of the tumors are often derived from the BCC tumors as both regions share somatic *PTCH1* mutations (Chiang et al., 2019). Using the same approach, we mechanically isolated tumor regions from histologically distinct BCC- and SCC-like regions of the Gorlin tumor for WES (Figure 5a). Samples were taken from the initial excision, which displayed the array of BCC- and SCC-like phenotypes (Region I, Region II, and Region V) (Figure 5b).

**Figure 5:**
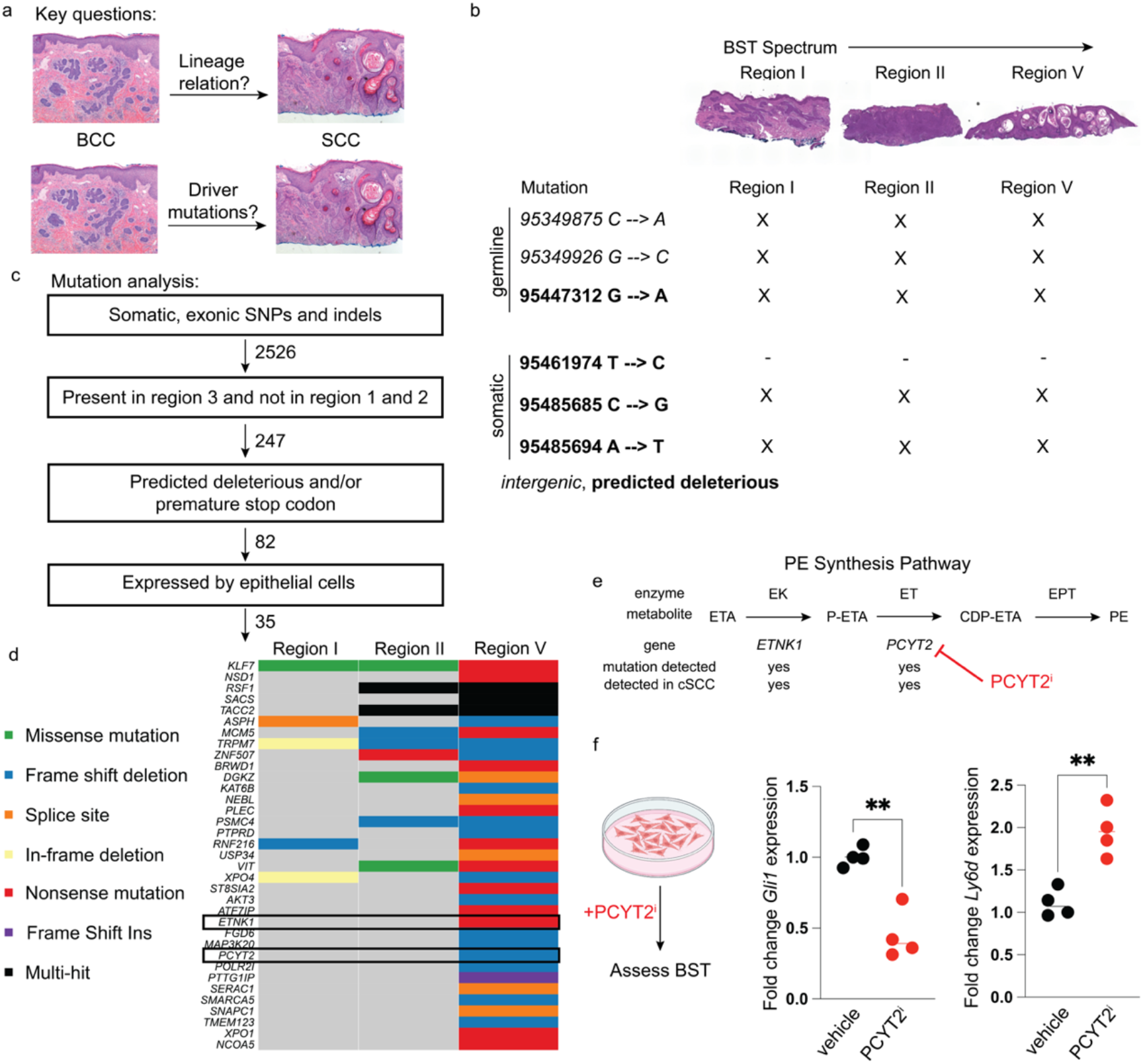
Mutational drivers of BST in resistant Gorlin tumors. a. Diagram of key remaining questions involved with the lineage relation and key driver mutations between the BCC- and SCC-like regions within the tumor. b. Diagram of tissue utilized for WES showing detailed diagrams of where the tissue came from. c. Key filtering steps used for identifying candidate somatic mutations. d. Germline and somatic mutation analysis of *PTCH1* from Excision-1 and Exision-2. e. Summary of the filtered 35 mutations focused on Regions 1-3. f. Gene expression of Gli1 and Ly6d from ASZs that were treated with PCYT2 inhibitor analyzed using student’s t-test with Welch’s correction (p=0.0047 and 0.0044)

To address the question of lineage, we found that the germline *PTCH1* mutations are shared across all regions of both tumors confirming their origins from the same patient (Figure 5b). Importantly, somatic mutation analysis revealed that there are predicted deleterious, acquired mutations at cytosine 95485685 and adenine 95485694 in all the regions sampled across the BST spectrum (Figure 5b and Supplementary Table S4). This would suggest that like BSCs, the SCC-like region of this tumor sample does carry the same *PTCH1* mutation as the BCC-like regions, indicating a lineage relationship between the two regions. Overall, this lineage analysis suggests the presence of one main tumor that spans the BST spectrum and an additional BCC-like tumor that is in proximity.

By WES, we were able to identify variants from each of the samples, which we were then able to classify as a particular variant subtype such as frameshift deletions, missense mutations, or splice site variants (Supplemental Figure S5c). As BCCs are generally diploid, most of the variants were either SNPs or deletions rather than copy number variations (Supplemental Figure S5c). Overall, we were able to identify 13234 somatic, exonic SNPs or indels with 247 being present in Region V but not Region I or II (Figure 5c). Of those 247, we found 82 of those were predicted to be deleterious and/or predicted to cause a premature stop codon (Figure 5c and Supplementary Table S5). Of those 82, we found that 35 were expressed by epithelial cells (Figure 5c, Supplemental Figure S4g, and Supplementary Table S6).

### Acquisition of additional mutations in PE synthesis pathway drives Gorlin-associated BST

Given the presence of BCC- and SCC-like tumors within a single tumor, we wondered which genetic drivers might induce a BST switch. Towards this end, we performed spatial InferCNV analysis to understand the predicted clonal relationships between the SCC- and BCC-like regions (Figure S4c-e) (Erickson et al., 2022). This analysis predicted that the two regions were clonally distinct with some predominantly BCC-like clones occupying adjacent regions to the SCC-like clones (Supplemental Figure 5e). Comparison with WES data did not reveal any obvious candidates in BST transition among genes with inferred copy number variation between the two regions, indicating that the mutational drivers of BST did not detectably alter copy number.

Using our WES data, we next focused on somatic mutations present only in the SCC-like region (Region V) and not present in the BCC-like regions (Regions I and II) (Figure 5b and c). Referencing the candidates from BSCs, we found a truncation mutation in *PTPRD* within the SCC-like (Region V) but not the BCC-like (Region I or II) areas (Figure 5e). Additional premature stop mutations that followed a similar pattern (detected in Region V but not Region I or II) were detected in 35 genes including *ETNK1*, *PCYT2*, and *NSD1*, representing novel candidate BST- driver mutations (Figure 5e and Supplemental Figure 4f and g). Of great interest, *ETNK1* and *PCYT2* encode sequential enzymes in the phosphatidylethanolamine (PE) synthesis pathway, which are critical to the synthesis of PE from ethanolamine (Patel and Witt, 2017). Mutations in *ETNK1* and *PCYT2* have been detected in cSCC (Figure 4e). We next measured whether the inhibition of the PE synthesis pathway could drive BST. Using PCYT2 inhibitors (PCYT2^i^) on our ASZ mouse BCC cell line, we found that treatment both reduced *Gli1* expression and increased *Ly6d* expression, consistent with a transition from an HH-dependent BCC-like to an HH-independent SCC-like state (Figure 5e) (Gohil et al., 2013). Overall, these observations suggest that mutations within the PE synthesis pathway may be unrecognized drivers of BST.

## DISCUSSION

In this work, we comprehensively profile a SMO^i^ suppression-resistant tumor arising within a patient effectively treated for 10 years and demonstrate the acquisition of resistance through BST in part through driver mutations in the PE biosynthetic pathway. The creation of SMO^i^ changed the standard of care for patients with basal cell nevus syndrome, providing a non-surgical option to suppress tumor growth with few escapers (Tang *et al.*, 2012). A central question was whether *LY6D* persister populations could grow out as SCC-like populations as seen in sporadic BCCs (Ransohoff et al., 2015). Analysis of the resistant tumor showed clear SCC-like histological features, raising the possibility that BCC tumors from Gorlin patients, like sporadic BCC tumors, can undergo BST as a resistance mechanism leading to a full outgrowth of an SCC-like tumor. From a clinical perspective, this is an important observation as it indicates that Gorlin patients that utilize SMO^i^ as a non-surgical option to reduce tumor burden are also susceptible to developing resistant tumors. Access to this clinical sample provides a unique opportunity to comprehensively profile and deconvolute the mechanisms at play driving resistance.

Using a combination of scRNA-Seq and 10x spatial transcriptomics, we were able to comprehensively profile this tumor and found that there are clear BCC- and SCC-like populations of cells that are spatially localized within the tumor. With this array of BST phenotypes, we were not only able to better understand the various resistance mechanisms associated with this case, but we were also able to further our understanding of BST as a general resistance mechanism. Using WES, we were able to show that the BCC- and SCC-like portions within the tumor are related by lineage and occupied adjacent yet distinct regions within the tumor. Using a combination of scRNA-Seq datasets from nBCC, rBCCs, and SCCs, we defined the BST spectrum and subsequently placed the different Gorlin tumor epithelial populations along the spectrum of BST. We found that the SCC-like portion of the Gorlin tumor was most like rBCC tumors that displayed extensive BST.

Additionally, the relatively low mutational burden in Gorlin BCCs compared to sporadic BCCs provided a unique opportunity to investigate novel genetic drivers of BST. Among the mutations detected in the SCC-like portions of the tumor, but not the BCC-like portions of the tumor are two genes encoding two enzymes in the phosphatidylethanolamine (PE) pathway. Interestingly, ethanolamine and PE are used interchangeably in human keratinocyte culture suggesting an essential role for PE in the maintenance of basal keratinocyte identity. Disruption of this pathway might then be expected to result in keratinocyte differentiation (Tsao et al., 1982). Our functional data suggest that inhibition of PE synthesis leads to BST in vitro, indicating that these mutations may contribute to BST transition in this tumor.

It is interesting to speculate as to why more resistant cases have not been seen with Gorlin patients on SMO^i^ as compared to sporadic BCCs. It is possible that this could be explained by the vast difference in mutational burden between sporadic BCCs and Gorlin tumors, where there are far fewer mutations seen in Gorlin tumors. In this instance, it took 10 years to develop resistance to SMO^i^ raising the possibility that given enough time, other Gorlin patients will develop resistance to SMO^i^ through the acquisition of additional mutations.

## MATERIALS & METHODS

### Human Samples

Patient samples were obtained through written informed consent and de-identified. All protocols for sample acquisition and usage are reviewed by the Stanford University Institutional Review Board, protocol #18325 (Stanford, CA).

### Human Sample Processing and Isolation

Human tumor samples were briefly rinsed with 1x PBS before being chopped and minced into pieces less than 1 mm in diameter. After mincing, tumor pieces are transferred to a 50-mL conical tube and 40 mL of 0.5% collagenase solution (Gibco; 17-100-017) in DKFSM media (Gibco; 10744-019). The minced tumors are then incubated at 37°C with rotation for 2 hours. At the end of the 2-hour incubation, 5 mL of 0.25% Trypsin (Gibco; 25200056) was then added to the 50-mL conical tube, which was then further incubated at 37°C with rotation for an additional 15 minutes. 5 mL of FBS was then added to the cellular suspension.

### scRNA-Seq Library Preparation and Sequencing

As per the directions of the Chromium Single Cell 3ʹ Reagents Kit v2 (following the CG00052 Rev B. user guide), mouse live FACS-sorted cells were washed in PBS containing 0.04% BSA and resuspended at a concentration of approximately 1,000 cell/μL. We did two replicates of the same sample to capture large amounts of cells for extensive analysis. We aimed to capture 10,000 cells for each replicate. Each library was sequenced on the Illumina NovaSeq platform to achieve an average of roughly 30,000 reads per cell.

### scRNA-Seq Library Processing, Quality Control, and Clustering

FASTQ files were aligned using 10x Genomic Cell Ranger 3.1.0 and aligned to an indexed hg38. Standard workflow using Seurat v3 with quality control parameters of 200-5000 gene detection and mitochondrial percentage under 10% were utilized to filter cells. Replicates were merged for analysis. Cells were presented in two-dimensional space using UMAP with the following sample-specific parameters and subset workflow listed below. For the Gorlin sample, cells were clustered using the top 10 PCs with a resolution of 0.3. Based on marker gene expression of epithelial markers (*KRT14*) and tumor-specific markers (*GLI1*), we were able to subset a tumor-specific population for the normal epithelium. For subsequent analysis, tumor epithelial cells were subsetted and then re-clustered using the top 10 PCs with a resolution of 0.3.

### scRNA-Seq Analysis

Marker genes were found for each cluster using the standard Seurat pipeline and parameters such as log(fold-change) > 0.25 as a cutoff for performing differential gene expression. For gene scoring, we used the AddModuleScore function built into Seurat. Gene Ontology analysis was conducted using marker genes for the various clusters, which were submitted to MSigDB collection from the Broad Institute. For Monocle-based lineage analysis, clusters and variable genes information is pulled from mouse-SCAM Seurat object and taken as input to construct pseudo time trajectory using the semi-supervised Monocle 2 (2.14.0) workflow.

### RNAScope

All RNAScope experiments were performed using the Multiplex Fluorescent v2 system (ACD; 323100) as per manufacturer protocols. Briefly, human and mouse tumors were fixed with 4% paraformaldehyde overnight at 4°C, paraffin-embedded, and sectioned at 5 μm. We followed the formalin-fixed, paraffin-embedded (FFPE) protocol as per manufacturing instructions. The following human probes were used: Hs-LY6D (ACD; 484681-C1) and Hs-*GLI1* (ACD; 310991-C2). For protein staining, we used the chicken K14 antibody (1:500; BioLegend SIG-3476-100) with the anti-chicken Alexa647 (1:500; Invitrogen; A-21449) secondary antibody. Nuclei visualization was with Hoechst 33342 (1:1000; ThermoFisher; 62249). Imaging was conducted using a Lecia SP8.

### Visium Sample Preparation, Library Preparation, and Sequencing

Human tumor samples were fixed in 4% PFA overnight, paraffin-embedded, and sectioned at 7μm. Each Visium capture area contained distinct regions of the same tumor sample. Visium slides were deparaffinized, stained, imaged, and decrosslinked according to 10x genomics demonstrated protocol (10x protocol, CG000241). H&E stained tissue was imaged using a Keyence microscope in accordance with 10x genomics Visium protocol. Libraries were constructed as described in 10x user guide and sequenced on the Illumina NovaSeq 6000 system S1-200 flowcell (10x protocol, CG000239).

### Visium Library Processing, Quality Control, and Clustering

Visium data were preprocessed using Space Ranger (get version from SG) and mapped to the GRCh38 genome. Further analysis was performed in using the Seurat R package. We normalized UMI counts using the SCTransform feature and integrated the four capture areas using anchors before performing dimensionality reduction by PCA. The FindClusters function was applied to assign identities on the basis of shared gene expression.

### Spatial transcriptomic analysis

Merged Visium data from the four capture areas was scored using the SpatialFeaturePlot feature in the Spatial Seurat R package. Spatial transcriptomic data was integrated with two scRNA-Seq datasets from the same sample using the SCTransform feature. After transferring with anchors, the gene signatures associated with each annotated cell type from the scRNA-seq data was used to score clusters from the spatial dimensionality reduction. These were manually examined to assign descriptive names to the clusters.

### CellTrek

CellTrek was used to deconvolute spot identity from the spatial transcriptomic data (Wei *et al.*, 2022). The data were co-embedded using *traint* and charted using default settings in the CellTrek package. Images were exported from the accompanying visualization tool.

### *In vitro* BST assay

ASZ_001 cells (known as ASZs) were cultured as previously described in M154CF (ThermoFisher; M154CF500) with 2% chelated FBS, 1% penicillin/streptomycin (Gibco; 15140122), and 0.05 mM CaCl2 (Haensel *et al.*, 2022). They were treated with vehicle (DMSO) or 1uM PCYT2 inhibitor (meclizine) for 48 hours after which RNA was isolated using TRIzol Reagent (Invitrogen; 15596026) and QIAGEN Rneasy Plus kit (QIAGEN, 74134). Gene expression analysis was performed using TaqMan RT-PCR Mix (Applied Biosystems, 4392587) was used for reverse transcription and subsequent quantitative PCR. Each reaction was performed in duplicate and values were normalized to *Gapdh* using the following probes/primers: Mm00494654, Mm00521959, Mm99999915 (Applied Biosystems).

### InferCNV

Spatial CNV analysis was performed using the SpatialInferCNV R package (Erickson *et al.*, 2022). We started by manually annotating the epithelial spots of Visium slide (IV) either as ‘Normal’, ‘BCC-like’, or ‘SCC-like in loupe browser based on histological examination. We assigned the ‘Normal’ spots as a reference set and ran the inferCNV program to generate a heatmap indicating copy number predictions with an accompanying dendrogram of probable lineage relationships. We manually selected dendrogram nodes, assigned them clone names, and exported the clonal predictions to loupe browser to spatially map inferred copy number variation.

### Whole Exome Sequencing and Analysis

Samples were selected for WES based on their unique histological features- one nodular BCC-like, one infiltrative BCC-like, and one SCC-like in appearance from the tumor biopsy and three nodular BCC-like regions from the re-excision sample. FFPE tissue was collected by microtome and processed using the Agilent FFPE DNA library kit (Agilent; 5191-4079) in order to achieve at least 100ng gDNA. The raw reads were aligned to the human hg38 reference genome using the BWAMEM algorithm, the resulting BAM files undergo processing using GATK4. Initially, the mapped reads were sorted and PCR duplicates, which may arise during library preparation and sequencing, were identified and marked. Base quality score recalibration (BQSR) was then performed using GATK4’s recommended human variant sites to enhance the accuracy of base calls. SNPs and Indels were called from the processed BAM files using HaplotypeCaller and Mutect2 variant-calling algorithms. Variants called were annotated using Variant Effect Predictor (VEP) tool. MAF files generated from the VCF files were used for visualization using maftools.

## DATA AVAILABILITY STATEMENT

The human-specific sequencing generated from this study is being deposited in the dbGaP database and will be made available at the time of publication. Previous human BCC scRNA-Seq data used in this study are available in dbGAP database under the accession code <u>phs003103.v1.p1</u>. The resistant human BCC scRNA-Seq publicly available data used in this study are available in the NCBI Gene Expression Omnibus under the accession code: <u>GSE123814</u>. The human SCC scRNA-Seq publicly available data used in this study are available in the NCBI Gene Expression Omnibus under the accession code: <u>GSE144240</u>.

## CONFLICT OF INTEREST

The authors state no conflict of interest.

## ACKNOWLEDGEMENTS

Cell sorting for this project was done on instruments in the Stanford Shared FACS Facility. The following NIH Shared Instrument Grants were used to fund equipment used in the Stanford Shared FACS Facility: NIH S10RR025518-01 and NIH S10RR027431-01. Imaging was conducted at the Stanford University Cell Sciences Imaging Core Facility. The following NIH Shared Instrument Grant was used to fund equipment used in the Stanford University Cell Sciences Imaging Core Facility: NIH 1S10OD010580. Visium processing and sequencing was carried out at the UC Irvine Genomics, Research, & Technology Hub, which is supported by P30CA-062203. The work is funded by the following grants: A.R.J is supported by NIH T32AR007422-40; D.H. is supported by NIH 5T32AR7422-37 and NIH 1F32CA254434; A.E.O. is supported by NIH 1R01AR04786 and NIH 2R37ARO54780. The diagrams shown in Figures 1a and 5f were created with BioRender.com.

## AUTHOR CONTRIBUTIONS

A.R.J., D.H., and A.E.O conceived, designed, and interpreted experiments. A.R.J., D.H., and A.E.O wrote the manuscript. A.R.J. and D.H. executed all wet lab-based experiments. A.R.J., D.H., and S.G. conducted all computational experiments. All authors revised the manuscript.

## TABLES

**Supplementary Table S1**: Marker genes from the scRNA-Seq of the clusters from the human Gorlin sample (Figure 2a).

**Supplementary Table S2**: Marker genes from the Visium spatial Seurat object in (Figure 2d).

**Supplementary Table S3**: Marker genes from the scRNA-Seq of the clusters from the human Gorlin tumor epithelium (Figure 3b).

**Supplementary Table S4**: WES analysis of *PTCH1* from the different tumor regions (Figure 5b).

**Supplementary Table S5**: Global WES analysis from the different tumor regions (Figure 5c).

**Supplementary Table S6**: WES analysis of the key 35 genes from the different tumor regions (Figure 5d).

---

**Supplementary Figure 1:**
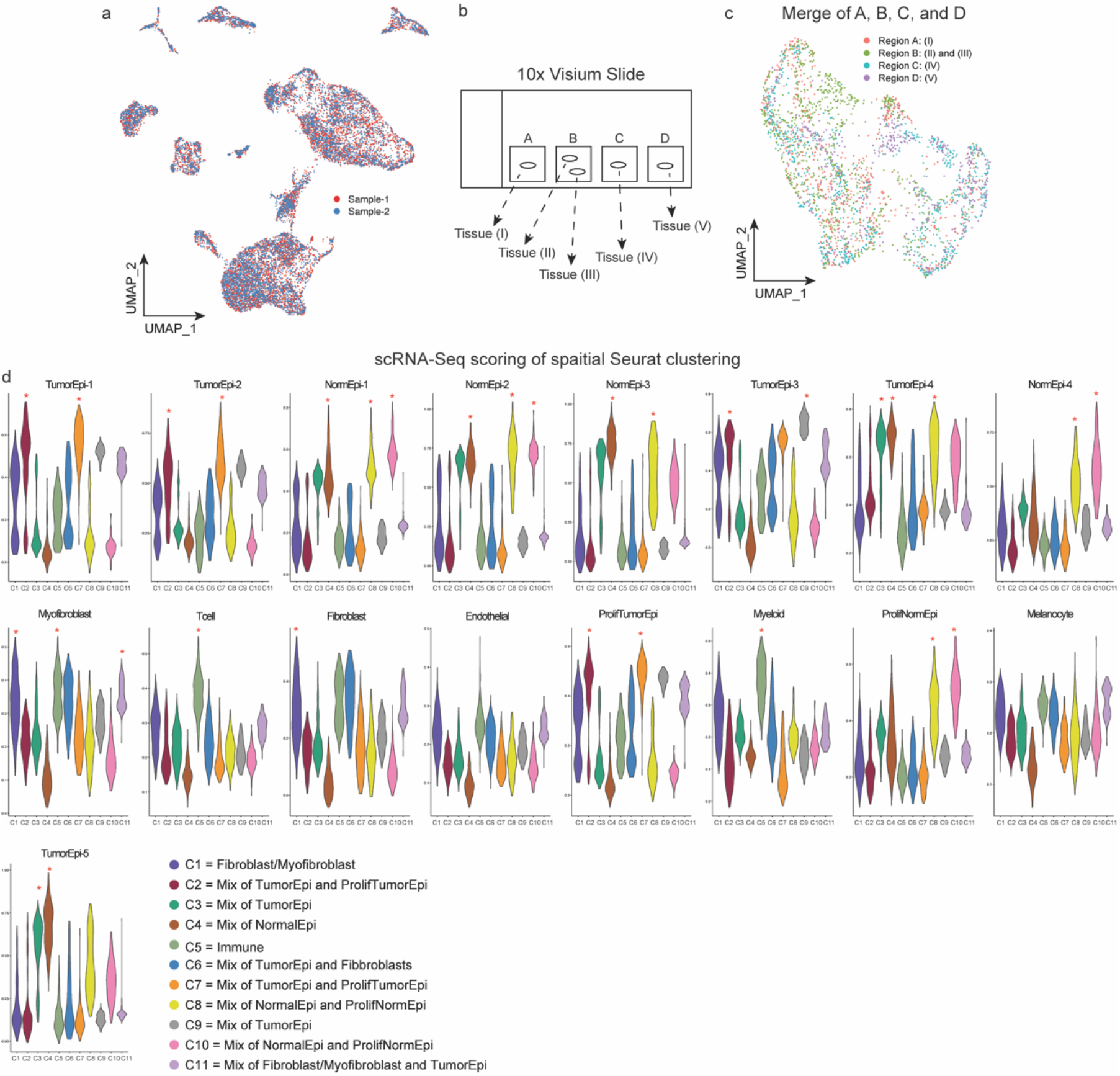
Coupled scRNA-Seq and spatial transcriptomics to profile resistant Gorlin tumor. a. UMAP plot of scRNA-Seq data. Two samples from the same patient were run through the 10x platform, sequenced, and then merged together. b. Diagram of the 10x Visium slide showing the placement of the various pieces of tissue in the different regions. c. UMAP plot of the merged spatial capture spots from the five different sections. d. Scoring of the different scRNA-Seq populations to identify the Visium populations asterisks indicate assigned identities.

**Supplementary Figure 2:**
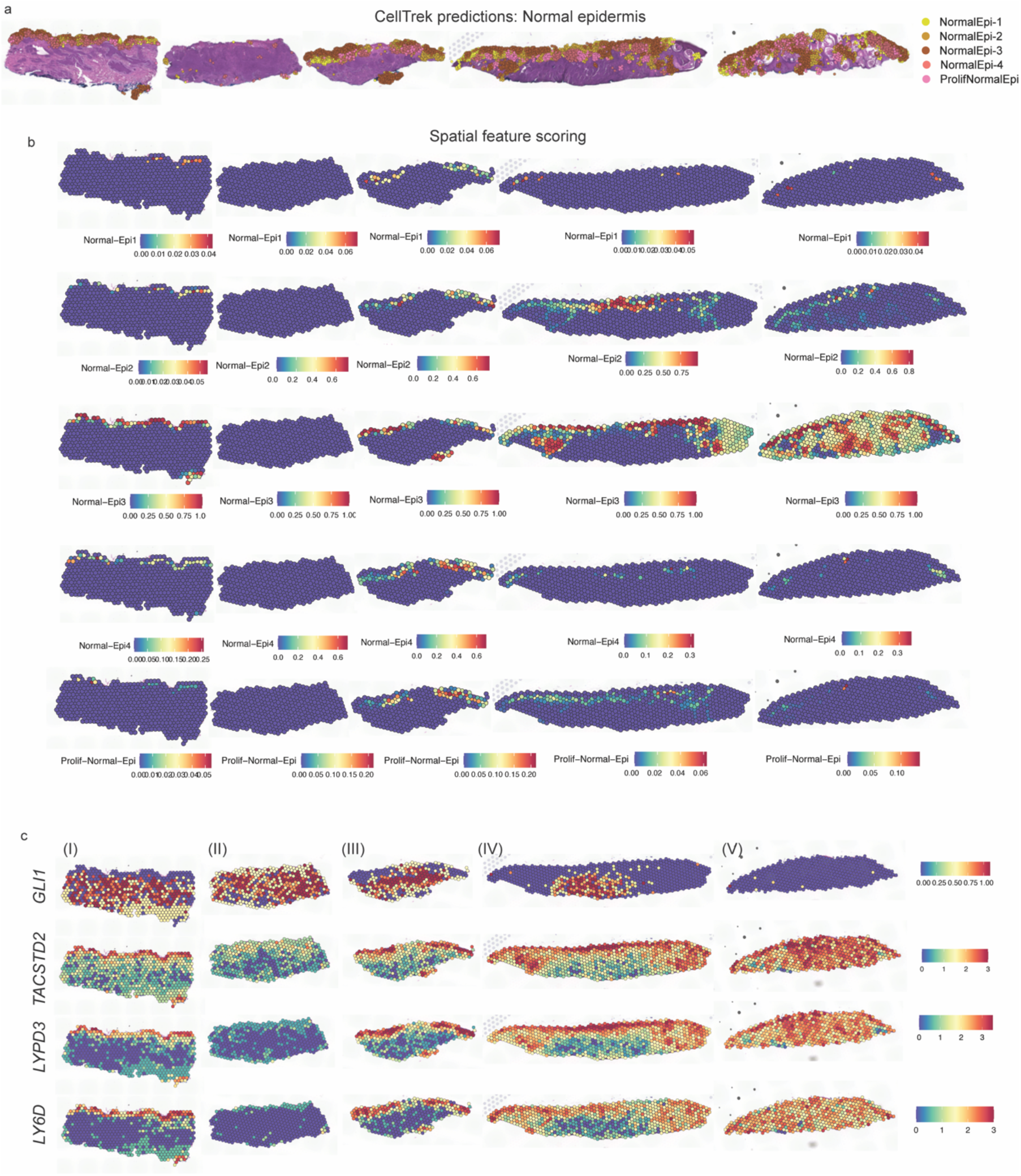
Spatial analysis and characterization of Gorlin tumor epithelial populations. a. CellTrek projection of the normal epidermal clusters onto the Visium analysis. b. SpatialSeurat cluster scoring for the normal epidermal populations on all tissue sections. c. Spatial feature plots of *GLI1*, *TACSTD2*, *LYPD3*, and *LY6D* for all tissue sections.

**Supplementary Figure 3:**
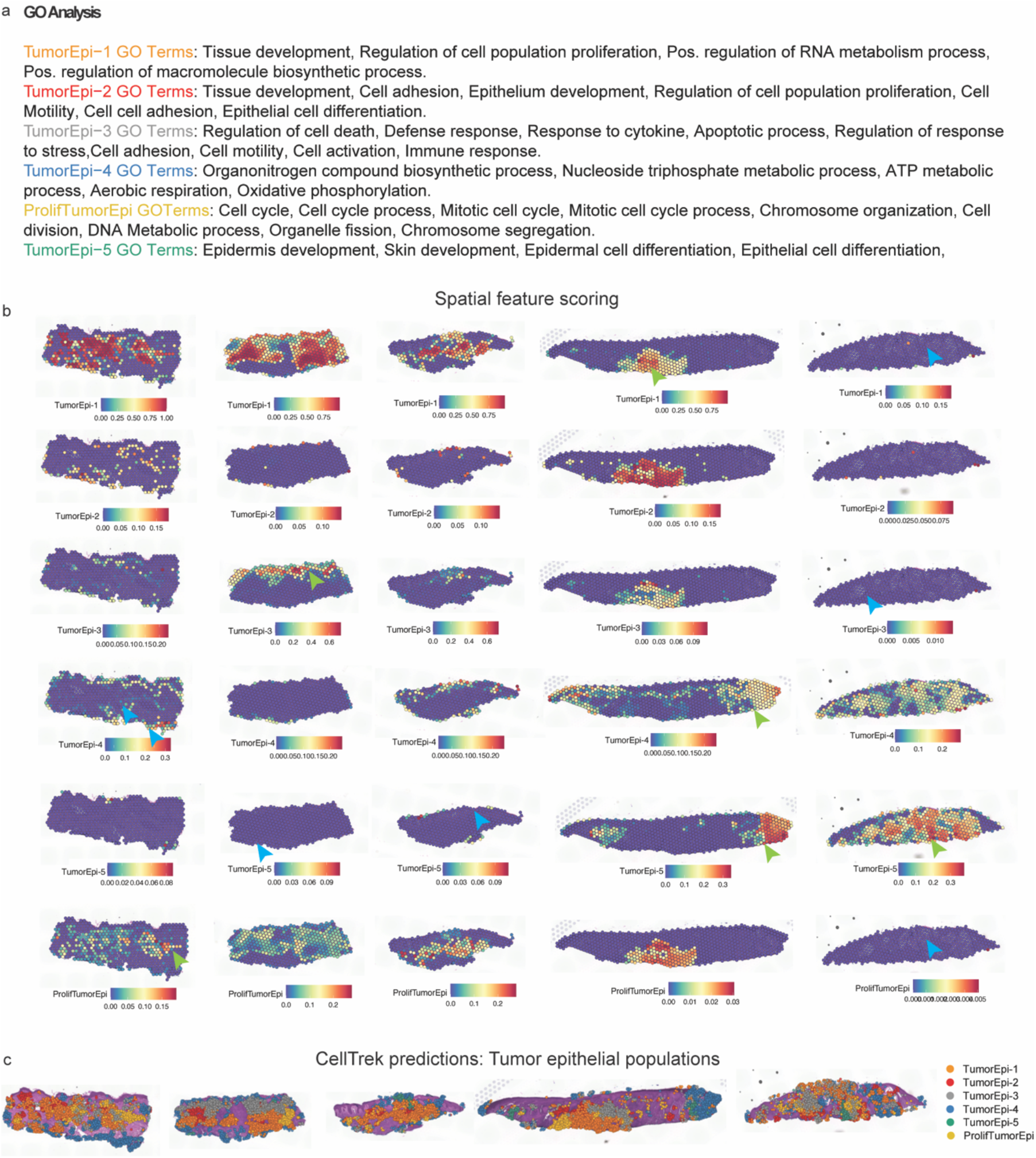
Spatial analysis and characterization of Gorlin tumor epithelial populations. a. GO analysis of the different tumor epithelial populations. b. Spatial gene scoring for the tumor epithelial populations on all the various tissue sections. c. CellTrek projection of the tumor epithelial clusters onto the Visium analysis.

**Supplementary Figure 4:**
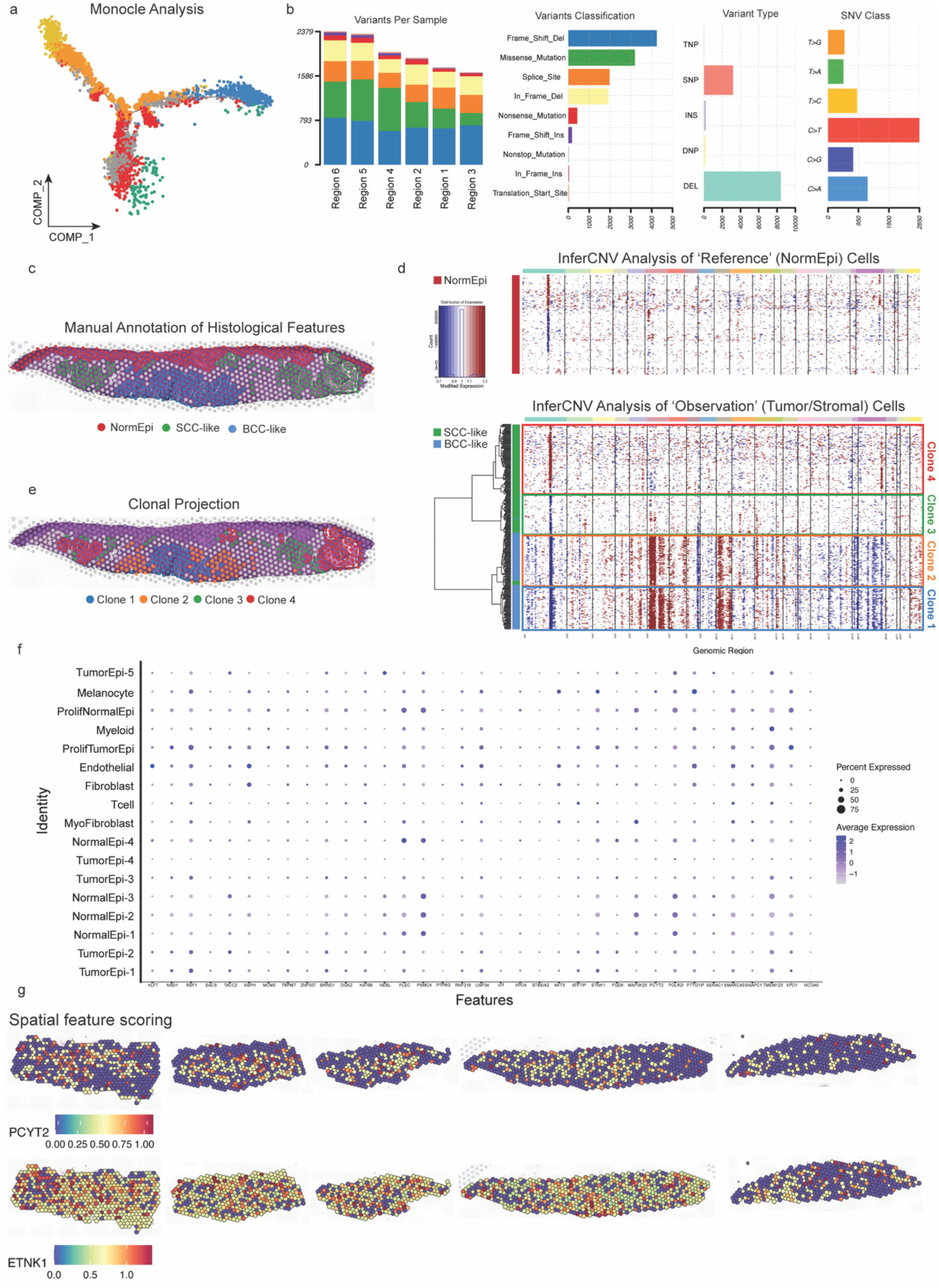
WES analysis. a. Monocle projection of the tumor epithelial clusters with the same cluster and color information in Figure 3b. b. WES analysis summary; From left to right: Summary of variants found from each sample; Variant classification summary from all samples; Variant type from all samples; SNV class from all samples. c. Dot plot showing gene expression of all 35 candidates which have epithelial expression. d. Spatial feature plots of *PCYT2* and *ETNK1* for all tissue sections.

## Notes

### Competing Interest Statement

The authors have declared no competing interest.

